# Intraspecific plant-soil feedbacks alter root traits in a perennial grass

**DOI:** 10.1101/2025.03.11.642669

**Authors:** Ceyda Kural-Rendon, Natalie E. Ford, Kara Hooser, Maggie R. Wagner

**Affiliations:** University of Kansas; Pennsylvania State University

## Abstract

Drought is a common stressor faced by plants and their associated microbiomes. Projected climate data point toward an increase in the severity and frequency of extreme precipitation events, such as drought. Previous research has shown that long-term exposure to drought can shape plants’ genomes, resulting in genetic variation for drought tolerance. We hypothesized that these genetic changes also affect patterns of microbial colonization in the rhizosphere, potentially feeding back to influence plant drought responses. Here, we tested 33 rhizosphere soils conditioned by 33 genotypes of *Tripsacum dactyloides* (eastern gamagrass) that originated from native populations across a precipitation gradient in the southern plains of the United States. We used these 33 rhizosphere soils as inocula for a fully factorial experiment to test the responses of conspecific plants to the differentially conditioned soils under drought or well-watered conditions. Variation in aboveground traits such as shoot length, weight, and root-to-shoot ratios was primarily explained by watering treatment. However, many belowground traits, such as root anatomical and architectural traits, were more likely to be affected by the genotype of the conditioning plant. Of the traits we measured, only aerenchyma area was affected by the interaction between current watering treatment and genotype of the conditioning plant. Ultimately, both the current watering treatment and conditioning plant genotype altered plant physiological traits and the associated microbiome. The differential intraspecies plant-soil feedback dynamics driven by plant local adaptation will be key to understanding future plants’ responses to rapidly shifting climates, in both restoration projects and agricultural systems.

## Introduction

The plant-associated microbiome is an important contributor to a plant’s ability to adapt to its environment. Arbuscular mycorrhizal fungi (AMF), for example, are thought to have assisted plants in their transition from solely aquatic ecosystems to dry land (Taylor et al. 1995). Plants are able to shape their associated microbial communities through functional traits such as root hair density, a key feature of the rhizosphere microbial habitat, and root exudates, which contain diverse sugars and phytochemicals that influence microbial growth (Ma et al. 2022, Wang et al. 2024). Many of these plant functional traits are genetically determined. For example, the ability to produce certain exudates that can encourage or deter the growth and association of some microbes and not others are a combination of genetic traits that—if beneficial to fitness—can become fixed in a population over time (Rolfe et al. 2019, Pacheco-Moreno et al. 2024). Thus, a plant’s associated microbiome can be seen as an “extended phenotype”, as it is modifiable by the plant’s phenotypic expression of root exudates and other root traits that affect microbial colonization (Wagner 2021).

Plants interact with the microbes in soil over time, recruiting some microbes that can then grow inside the plant’s roots (root endosphere) and on and around the plant’s roots (rhizoplane and rhizosphere, respectively) (Lundberg et al. 2012). For instance, a plant affects the composition of the soil microbiome in its immediate vicinity (the rhizosphere), which forms the surrounding microbial community that has the opportunity to colonize the endosphere (Hartman and Tringe 2019). The influence of plant genotype and phenotype over the microbiome may decline with distance from the root, with plants having the least amount of control over microbial communities in the surrounding bulk soil (Zhou et al. 2022), by nature of the concentration of root exudates and their ability to diffuse through the soil.

The process of plants differentially influencing the microbial communities in the surrounding soil, by deterring or promoting the growth of the constituent species, is called conditioning (Hu et al. 2018). A plant conditioning its rhizosphere microbes over time can lead to plant-soil feedback, in which both the plant and the microbes are interacting with each other in ways that alter the growth of both other plants and microbes in the vicinity (Bever et al. 1997, De Vries et al. 2023). Plant-soil feedback (PSF) is incredibly context-dependent and can be negative (when a plant conditions a microbiome that benefits competing plants) or positive (when a plant conditions a microbiome that benefits itself) depending on the species of the conditioning plant, the surrounding plants, the microbial communities, environmental factors, and many other biotic and abiotic factors (Smith-Ramesh and Reynolds 2017). However, all types of PSF begin with genetically-controlled plant traits that alter the rhizosphere microbiome in a deterministic fashion.

The mechanisms by which microbiomes can feed back to influence plant fitness are diverse. Notably, microbes can affect the physical structure of plants, including root architecture and root anatomy (Calzavara et al. 2018, Grondin et al. 2024). These traits not only respond to and are shaped by the activity of surrounding biota (Gao et al. 2023), but are also important for the plant’s tolerance of water stress. Large aerenchyma and cortical cell size, for example, have been found to help plants withstand flooding and drought stress (Yamauchi et al. 2013, Chimungu et al. 2014, Yamauchi et al. 2021a). AMF specifically have been found to increase root length, diameter and surface area in *Panicum virgatum* under water-stressed conditions (Basyal and Emery 2021).

A changing environment can therefore affect how a plant conditions its associated soil microbes, and how future plants will respond to soils conditioned by plants that may have experienced a different climate. One important shifting environmental factor for plants and their associated microbiome is precipitation. Severe precipitation events are projected to become more frequent, whether that be drought or flooding events (Trenberth 2011). The severity of these challenges will differ for drought-adapted *vs.* non-adapted plants. Genotypes adapted to historically drier conditions tend to have higher water use efficiency compared to genotypes of the same species that are adapted to historically wetter sites (Dudley 1996, Knight et al. 2006). Plants along the drier side of an aridity gradient have been found to allocate more resources to root development, which can increase water acquisition under arid conditions (Welles and Funk 2021). Plants that are better adapted to drought conditions are likely to remain relatively phenotypically stable during drought periods, which may translate into stability of plant-soil feedback dynamics (Van Der Putten et al. 2016). While the roles of plant population genetics and local adaptation in shaping plant performance under drought stress are well described, the influence of plant soil feedback on microbial recruitment—and, by extension, its impact on plant trait expression during drought—remains poorly understood.

Here, we use soils conditioned by 33 genotypes of *Tripcasum dactyloides* (eastern gamagrass) to test how the legacy of historic precipitation affects how plants shape their soil microbial communities, and how future conspecifics respond to those conditioned soils. *T. dactyloides* is a perennial bunchgrass native to most of the United States east of the Rocky Mountains. The range of this species coincides with a 10,000-year-old precipitation gradient, with the eastern sides of Texas, Oklahoma and Kansas receiving about 1270-1524 mm of precipitation annually, gradually decreasing to about 381-508 mm of precipitation annually on the western borders of those states (Axelrod 1985). Two recent studies have demonstrated that genetic variation among accessions of this species is partially explained by historical mean annual precipitation (MAP).

For instance, *T. dactyloides* genotypes originally from the western portion of this gradient have higher densities of fungal root endophytes than genotypes originally from the eastern portion of the gradient, suggesting that historic precipitation levels have shaped how this species forms its microbiome (Kural-Rendon et al. 2023). Furthermore, *T. dactyloides* genotypes from locations with greater mean annual precipitation are taller than genotypes from drier sites when grown in a common garden environment (Kiniry et al. 2023), suggesting that MAP has shaped genetic variation in this species. Finally, recent experimental work has shown that interactions with soil microbes can improve water-use efficiency of *T. dactyloides* during acute drought (Ginnan et al. 2024). Together, these results show that eastern gamagrass is a promising study system for exploring intraspecific genetic variation for plant-soil feedbacks under drought conditions.

To investigate how historic precipitation shapes genetic variation for plant-microbiome interactions, we compared rhizosphere soils that were conditioned by 33 genotypes of *T. dactyloides* growing side-by-side in a common garden. The original collection sites of these 33 genotypes span the southern plains precipitation gradient (**Figure 1**). To assess the relationship between historic precipitation and plant-soil feedbacks, we grew a common “tester” genotype of *T. dactyloides* in each of the 33 differentially-conditioned rhizosphere soils. Additionally, to test how current water availability may affect the phenotypic traits, as well as how these traits are shaped by the interaction between current water availability and historic precipitation, we exposed half of the experimental plants to a drought treatment, and the other half to a well-watered treatment. We used this experimental setup to investigate the following questions: (1) Do genotypes of *T. dactyloides* differentially condition their soil microbiota? (2) If so, does this genetic variation for rhizosphere conditioning correspond to historical patterns of precipitation at the genotypes’ home sites? (3) Does this differential conditioning affect how future conspecifics of *T. dactyloides* respond to drought?

**Figure 1.**
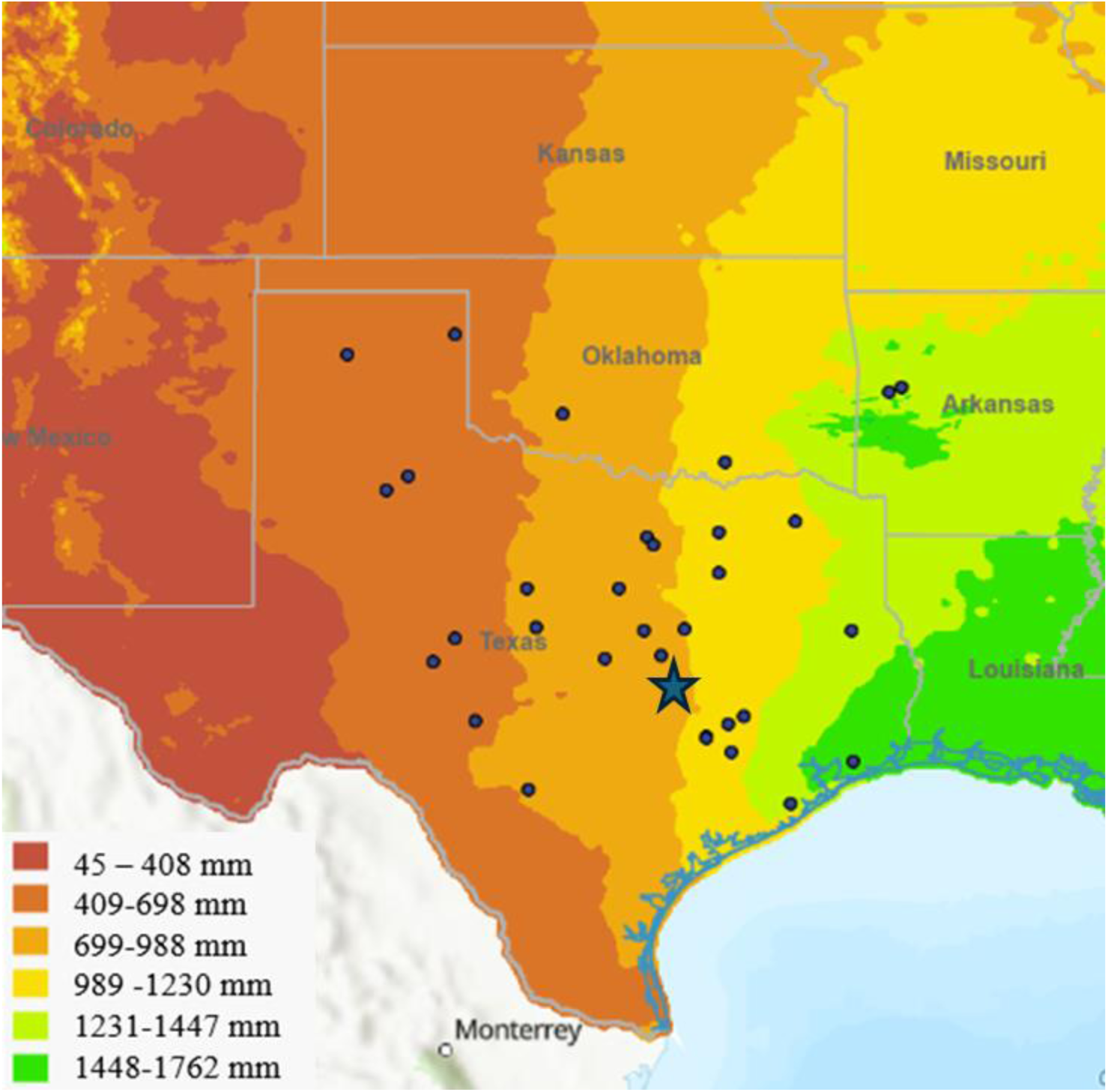
Map depicting the original collection sites of *T. dactyloides* genotypes, spanning an east to west precipitation gradient (mean annual precipitation shown in millimeters). The blue star denotes the common garden in Temple, Texas where collected genotypes (originally from blue points on map) grew for 4 years and where the resulting conditioned rhizosphere soils were collected. Map colors represent the mean annual precipitation.

## Methods

### Soil Inoculum Collections

The 33 focal genotypes of *Tripsacum dactyloides* were grown together in a randomized arrangement within a common garden at the USDA ARS Soil and Water Laboratory in Temple, Texas, USA for 4 years (**Figure 1**). In early March 2022 we collected loosened rhizosphere soil from about 6 inches under the base of each plant using a hand spade. For each genotype, we filled three 50 mL conical tubes with soil, disinfecting our trowel between genotypes. The soils were placed on ice until they were transferred to storage at -20 ℃ for future use as inocula.

We also sub-sampled these soils for amplicon sequencing by collecting 1.5 mL aliquots in Eppendorf tubes. We placed these sub-samples on ice until they were transferred to storage at - 20℃ for downstream DNA extractions.

### Experimental Design

In May 2022, we conducted a greenhouse experiment to assess the effects of the 33 collected rhizosphere soils on a common “tester” genotype. Seeds of the *T. dactyloides* cultivar “Pete” were soaked in a 3% hydrogen peroxide solution for 24 hours to increase germination (Gamagrass Seed Co., Falls City, Nebraska). They were sterilized in 75% ethanol and rinsed with sterilized water before planting.

The 33 rhizosphere soils conditioned by the 33 different genotypes of *T. dactyloides* were removed from their respective conical tubes and homogenized individually. Clean 256 cm^3^ pots were filled about three-quarters full with Berger BM7 potting soil that had been autoclaved twice at a 45-minute liquid cycle (Berger, Saint-Modeste QC Canada). A layer of one of the 33 soil inocula was added as 10% of the total volume (Lubin et al. 2021) to 8 total pots per inoculum. Once all 33 soils had been used to inoculate 8 pots each, 10 surface-sterilized “Pete” seeds were dropped on top of the inoculum layer in each pot, to account for the low germination rate of *T. dactyloides*. The rest of each pot was topped off with more sterile potting soil. Eight pots did not receive a soil inoculum and instead contained 100% sterile potting soil to serve as a control group. After germination, excess seedlings were removed promptly so that only one individual plant remained in each pot. Pairs of well-watered and drought plants that received the same inoculum (or no inoculum if they were in the control group) were randomized in trays and placed in the greenhouse, for a total of 4 pairs for each of the 33 soil inoculum types plus 4 pairs of plants in the un-inoculated control group, for a total of 272 plants (**Figure 2**). Trays of randomized plants were then numbered, and each tray or “plot” number is accounted for as a random effect in our statistical analyses to control for spatial variation in greenhouse conditions.

**Figure 2.**
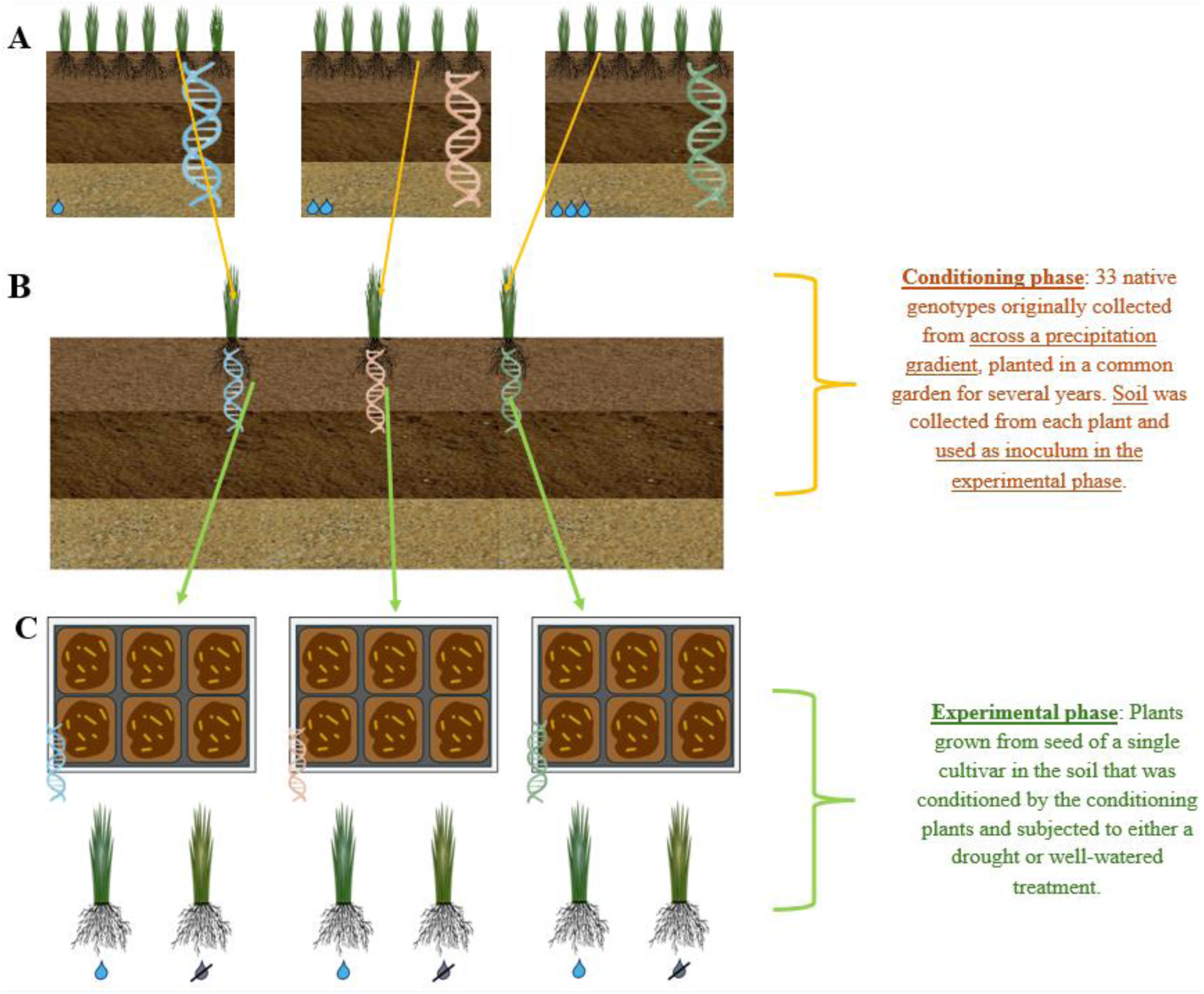
Schematic of experimental design. A. Three genotypes are depicted in their historical habitats, which differed in mean annual precipitation. Our study used a total of 33 genotypes spanning the Great Plains precipitation gradient. B. Each genotype was installed in a common garden in Temple, Texas, where they grew for 4 years prior to rhizosphere sampling. We hypothesized that during this “ conditioning phase”, phenotypic differences among the genotypes may have differentially shaped the surrounding rhizospheres. C. Rhizosphere samples collected from 33 genotypes in the common garden were used as inocula in the experimental phase. Each inoculum had a total of 8 replicates, 4 of which were given a drought treatment, and the other 4 the well - watered treatment.

Plants were allowed to grow for 2 weeks before the drought treatment and measuring began. During this initial phase, they were all watered every other day, or when the topsoil was dry. After the two-week acclimation period, beginning on May 19^th^, 2022, well-watered plants were given 100 mL of water twice a week, whereas droughted plants were given 50 mL of water once a week. Plant heights were measured weekly. Chlorophyll content was measured using a MC-100 Chlorophyll Concentration Meter (Apogee Instruments Inc., Logan UT) mid-experiment to confirm that drought treated plants were experiencing the effects of drought, which reduces chlorophyll content (Zhang et al. 2011). After 10 weeks, all watering was stopped for 3 days before roots and shoots were separated, weighed, and measured. Whole root systems were preserved in 75% ethanol for later root architecture, endophyte quantification, and anatomical analyses. Shoots were placed in brown paper bags and dried at 50 ℃ for 24 hours before being weighed.

### Root Architecture Scanning

Whole, intact root systems were cleaned of soil and then scanned in a plastic tray filled with deionized water using an EPSON V600 scanner (Epson, Los Alamitos, CA USA). Roots were manually separated and arranged to lie flat so that all root branches were visible. Using the RhizoVision software (Seethepalli et al. 2021) in whole root analysis mode, we converted the pixels to physical units at 600 dpi, and thresholded the images to level 200. We then selected the region of interest to exclude any border shadows that could have been mistaken as root. Root pruning was enabled in order to remove false positive root tips. The root pruning threshold was set to 5 pixels with two root diameter ranges: 0-0.30 mm and >0.30 mm. We then analyzed all images with the RhizoVision software to measure the average, median, and maximum root diameter; number of root tips; and root surface area for all root samples.

### Root Cross-Section Examination

Intact secondary lateral roots were selected from the preserved root systems. Three thin cross-sections of roots were taken about 5cm apart, beginning at but not including the root tip itself. The root sections were dyed using a 1.25% toluidine stain solution and placed on a microscope slide, with a glass cover placed on top. The root cross-sections were then viewed and photographed under the microscope at 4X magnification under the microscope (Accu-Scope 3000-LED, Accu-Scope INC., Commack, NY) using the attached Manual-AU-600-Excelis HD Camera.

All three root-cross section photos per plant (Figure 3) were analyzed using ImageJ (Schneider et al. 2012). The scale was set at 93.5 pixels per 100 microns. Images were cropped to focus on the root cross-sections, followed by a measurement of the area and perimeter using the polygon tool. To measure the combined area of the aerenchyma, we changed the image to 8-bit and used the wand tool to select the aerenchyma on the thresholded images. We used the circle tool to make selections around the metaxylem and calculate the combined area. The areas of aerenchyma and metaxylem were each standardized by dividing by the total root cross section area. The data from each of the three images from a single plant were then averaged before analysis.

**Figure 3.**
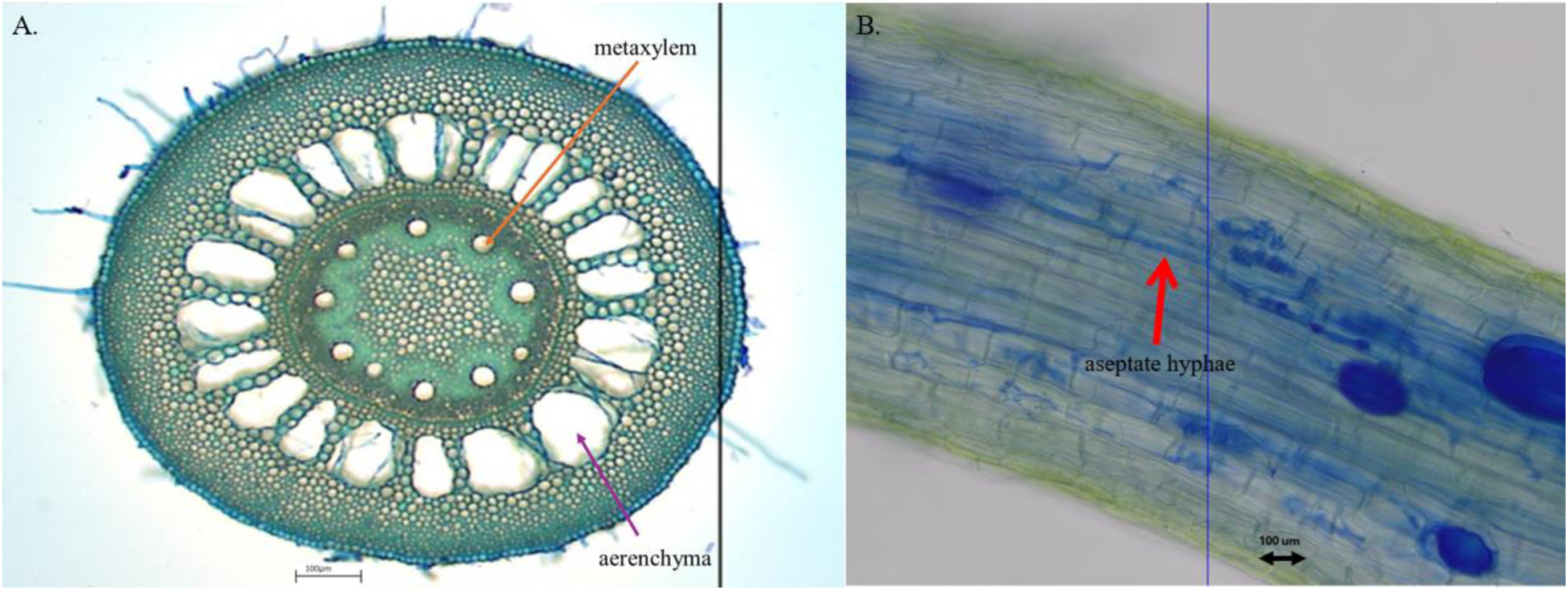
A. Cross section of a *Tripsacum dactyloides* root. Arrows indicate a singular aerenchym and a single metaxylem. B. Fungal endophytes in a T. dactyloides root, red arrow pointing out the aseptate fungi at 20 X magnification.

### Endophyte Quantification

The fungal endophyte cold-staining and quantification protocol was adapted from McGonigle et al. (McGONIGLE et al. 1990). Finer, secondary intact lateral roots that had been preserved in 75% ethanol were washed, cut into 1 cm long sections, and left in a 10% potassium hydroxide solution for 48 hours to clear plant tissue. Cleared roots were then transferred to a 1% hydrochloric acid solution for 24 hours followed by a 1% aniline blue staining solution for another 24 hours. Lastly, roots were left in a 14 parts 88% lactic acid, 1 part 100% glycerol and 1 part water destaining solution on a shaker at 30 rpm for 3-5 days. Destained roots were mounted on a microscope slide and covered with a drop of destain solution and a glass slide cover.

Fifty fields of vision from each sample were viewed at 20x magnification under the microscope (Accu-Scope 3000-LED, Accu-Scope INC., Commack, NY) using the attached Manual-AU-600-Excelis HD Camera. If fungal hyphae touched the microscope’s crosshair viewer at any point in the field of vision, the field was considered endophyte positive, and the type of fungi was recorded (septate or aseptate). Occasionally, both septate and aseptate hyphae were touching the crosshair viewed in the same field of vision, and these cases were recorded separately. If no fungal hyphae intersected the crosshairs viewed, the field of vision was considered endophyte negative. For each sample, we then calculated the total proportion of fields of vision that were endophyte positive, as well as specifically the proportion of fields of vision in which septate, aseptate or both were present. Oomycete presence in a sample was also noted, but densities were not quantified.

### DNA Extraction and PCR

To extract microbial DNA from the aliquots of the rhizosphere soils used as inocula, sterile garnet grinding material (BioSpec, Bartlesville, OK, USA) was added to every well of one 2.0 mL round bottom 96-well plate, followed by 250 mg of soil. We used 50 µL of the ZymoBIOMICS Microbial Community Standard (Zymo Research, Irvine, CA, USA) as a positive control and included a blank well as a negative control.

We then added 800 µL of lysis buffer (1M Tris pH 8.0, NaCl, ethylenediamine tetraacetic acid [EDTA]) to each well, followed by 20% sodium dodecyl sulfate. Samples were homogenized (20 Hz for 10 min; Ohaus HT Lysing Homogenizer, Parsippany, NJ, USA) and incubated at 55°C for 90 min. The lysate was removed from the garnet via centrifugation (6 min at 4500 x g) and transferred to a new 1.0 mL 96-well plate with 130 µL of 5 M potassium acetate. After an overnight incubation at -20°C, samples were thawed and centrifuged to remove protein complexes, then the supernatant was transferred to a new 1.0 mL round bottom 96-well plate with 1.5x volume solid-phase reversible immobilization (SPRI) bead solution (Cytiva Sera Mag SpeedBeads, polyethylene glycol 8000, NaCl, Tris-EDTA) and mixed thoroughly (Rohland and Reich 2012). After a 10 min incubation to allow DNA to bind to the SPRI beads, the sample plate was placed on a magnet rack (Invitrogen) for an additional 10 min. Once the SPRI beads were fully bound to the magnet, the supernatant was removed and two 80% ethanol washes were performed. All remaining ethanol was removed from the samples after the final wash, and the DNA was eluted from the magnetic beads with 37°C Tris-EDTA buffer, transferred to a sterile 0.45 µL V-bottom 96-well plate, and stored at -20°C until further use.

To amplify the bacterial 16S-v4 rRNA gene region, we used the 515f-806r primer pair (Parada et al. 2016). For fungal libraries, we used the ITS1f-ITS2 primer pair to amplify the ITS1 region of the rRNA gene (Smith and Peay 2014). For the 16S-v4 rRNA gene region, each PCR contained 0.4 µL of both the forward and reverse primers (at 10 µM), 5 µL of the 5Prime HotMasterMix, 0.5 µL of 10 mg/mL bovine serum albumin (BSA), 0.15 µL of 16S peptide nucleic acid (PNA) blockers (PNA Bio), 1.05 µL of PCR-grade water, and 2.5 µL of template DNA (Lundberg et al. 2013). The BSA was used to prevent non-specific binding of primers, while the PNA was employed to prevent binding of primers to 16S sequences in mitochondria and chloroplasts. Each amplifying PCR reaction for the ITS1 region included 0.4 µL of both the forward and reverse primers (at 10 µM), 0.4 µL of 10 mg/mL BSA, 4 µL of 5Prime HotMasterMix, 3.6 µL of PCR-grade water, and 1.2 µL of template DNA. To amplify the 16S-v4 gene region, we subjected reactions to an initial denaturation step of 95°C for 2 min, followed by 27 cycles of the following: an additional denaturation step of 95°C for 20 s, PNA annealing at 78°C for 5 s, primer annealing at 52°C for 20 s, and an extension step at 72°C for 50 s. A final extension step of 72°C for 10 min was also performed. To amplify the ITS1 region, we first denatured reactions at 94°C for 3 min, followed by 35 cycles of the following: an additional denaturation step at 94°C for 45 s, primer annealing at 55°C for 1 min, and an initial extension step of 72°C for 90 s. We followed this with a final extension step at 72°C for 10 min. A second, barcoding PCR was performed for both 16S and ITS amplicons, in which 2 µL of premixed barcoded Illumina forward and reverse adapters (at 10 µM in total), 12.5 µL of 5Primer HotMasterMix, 1.25 µL of 10 mg/mL BSA, 0.375 µL PNA, and 3.875 µL of PCR-grade water were mixed with 5 µL of template DNA. Samples were incubated for 2 min at 95°C as an initial denaturation step, followed by 8 cycles of: an additional denaturation step of 95°C for 20 s, a PNA annealing step at 78°C for 5 s, a primer annealing step of 52°C for 20 s, and an extension step at 72°C for 50 s. A final extension step was performed at 72°C for 10 min. PCR products were normalized and pooled using the Just-A-Plate kit (Charm BioTech, Cape Girardeau, MO, USA). Libraries were sequenced (2×250 bp) on an Illumina NovaSeq 6000 SP flow cell.

### Sequence Data Processing

We used cutadapt version 4.2 (Martin 2011) to process and trim paired bacterial and fungal reads simultaneously. Dada2 version 1.26.0 (Callahan et al. 2016) was then used to quality filter by truncating reads at the first instance of a quality score less than or equal to 2. We also discarded forward and reverse reads with expected errors higher than 2 basepairs, as well as sequences containing any ambiguous bases. Separately, 16S and ITS sequences were used to estimate error rates (using a minimum of 1e8 bases) for denoising. This was followed by dereplication of sequences into amplicon sequence variants (ASVs), merging pair-end reads, and removing chimeras. Taxonomy was assigned with the RDP training set (v19) for bacteria (Wang and Cole 2024) and the UNITE database (4 April 2024 release) for fungi (Abarenkov et al. 2024).

In R (version 4.3.2), ASVs assigned as mitochondria or chloroplast were omitted to remove plant contamination. We also removed ASVs that were unclassified at the Kingdom level, and any ASVs that were not present in at least 20 reads across 2 samples. This filtering process retained 1673 of the original 9063 bacterial ASVs (retaining 75.7% of all reads) and 242 of the original 1373 fungal ASVs (retaining 59.73% of all reads). The ALDEx2 package (version 1.34.0) (Gloor et al. 2016) was used to normalize ASV counts with a centered-log-ratio (CLR) transformation.

### Data Analysis

We used R (v 4.3.2.) for our analyses, along with packages phyloseq, vegan, genefilter, ALDEx2, tidyverse, lmerTest, and lme4 (Fernandes et al. 2013, 2014, Love et al. 2014, McMurdie and Holmes 2013, Gloor et al. 2016, R. Gentleman 2017, Wickham et al. 2019, Bates et al. 2015, Kuznetsova et al. 2017). All p-values were adjusted for multiple comparisons using the Benjamini and Hochberg correction (1995). We used raw counts to calculate the relative abundances of each taxon. Relative abundances of the top bacterial classes and families, and top fungal orders and genera are displayed in Supplemental Figures 1-4. Functional traits were assigned to fungal genera with the FungalTraits database (Põlme et al. 2020). Because we only collected one aliquot of conditioned soil per plant per genotype, we could not conduct statistical analyses to compare the microbiome community composition among genotypes. Instead, we used our data on fungal endophyte densities to assess genetic differences in soil conditioning.

**Figure 4.**
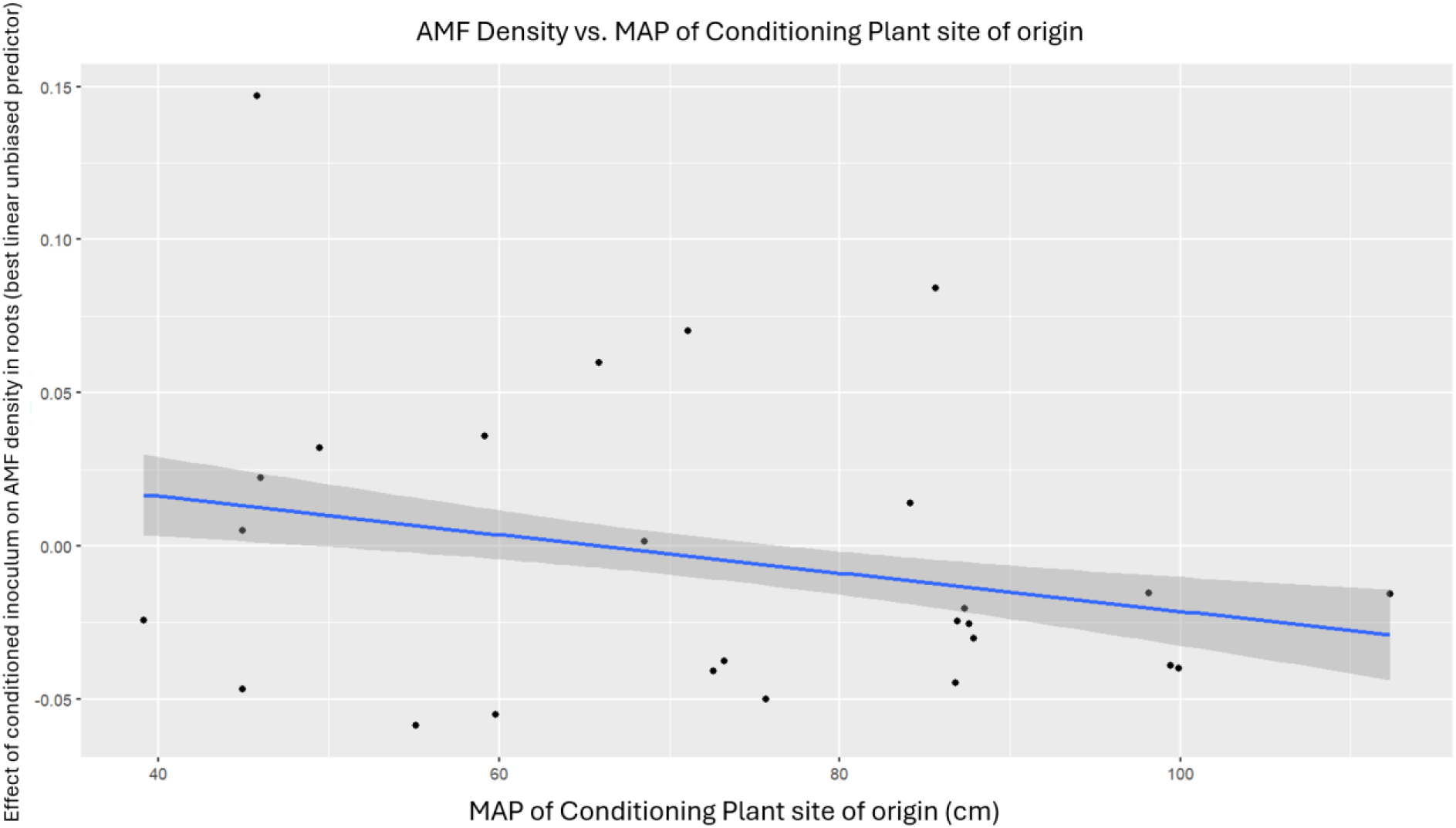
MAP of a conditioning plant site of origin was not correlated with AMF density (p >0 . 05, F=0 . 0118, df=1, 26, r =-0 . 037)

For our analyses of plant traits and endophyte densities, we used the following linear mixed model: *plant trait ∼ watering treatment +(1|plot)+ (1|conditioning plant genotype) + (1| treatment*conditioning plant genotype)*, where each plant trait was modelled separately. We treat the conditioning plant genotype as a random effect in our models, as the 33 genotypes used in this experiment were a random selection of the genotypes that exist along the precipitation gradient of interest. The final interaction term was included to determine whether the current watering treatment modified the influence of the conditioning plant’s genotype. We included the plot number where the plants were randomly arranged in pairs (one well-watered and one drought treated plant given the same inoculum) as a random effect in our models. Estimated marginal means (EMMs) were also calculated for each trait where watering treatment had a significant effect, in order to determine the treatment effect size. For traits for which conditioning plant genotype had a *P*-value<0.1 (as assessed by likelihood ratio test), we calculated the best linear unbiased predictor (BLUP) for each conditioning plant genotype.

To further discern whether the conditioning plant genotype effect was due to local adaptation of the conditioning plants to their home sites’ precipitation, BLUPs were then used as a response variable in a simple linear model with MAP of conditioning plant site of origin: *lm(BLUP ∼ MAP of conditioning plant’s site of origin)*. MAP of conditioning plant’s site of origin was the 30-year normal mean annual precipitation in millimeters of the site where the original conditioning plant was collected from wild populations of *T. dactyloides* (PRISM Climate Group). Statistical significance of this relationship was assessed using Spearman correlation. All p-values were adjusted using the Benjamini Hochberg correction (Benjamini and Hochberg 1995).

The plant phenotypic traits tested in our linear models were the area under the height growth curve of each plant, final root wet weight and height, final shoot height, wet weight, and dry weight. We did not include root dry weight because the roots were preserved in ethanol for other analyses and drying them could have compromised anatomical measurements. We also analyzed the root-to-shoot ratio based on both length and wet weight. For our root architecture analyses we used the following traits: average, median, and maximum root diameter, number of root tips, and root system surface area. Our root cross-section traits analyses focused on total root cross section area, aerenchyma area, and metaxylem area. For our endophyte quantification data we looked at overall endophyte density, septate hyphae density, and aseptate hyphae density.

Finally, our oomycete presence vs. absence data was analyzed with binomial logistic regression: *glmer(oomycete presence∼ watering treatment (drought or well-watered) +(1|plot) +(1|conditioning plant genotype), family =binomial)*. Finally, we tested whether several traits were correlated with fungal endophyte density, using a Pearson Correlation test: *endophyte density ∼ plant trait*.

## Results

### Variation in aboveground plant traits was attributable to watering treatment but not inoculum

The watering treatment predicted wet and dry shoot weight, shoot length, the total plant weight, wet root to shoot weight ratio and root to shoot length ratio (**Table 1**). Well-watered shoots were both 4.7 % taller and 28.3 % heavier (when wet and dry) than their droughted counterparts. Droughted plants had both higher root to shoot length (19.1% greater) and weight (13.2 % greater) ratios compared to well-watered plants. Watering treatment also predicted chlorophyll content mid-experiment, with well-watered plants having 13.9% higher chlorophyll density than droughted plants (**Table 1**). Watering treatment did not predict the area under the height growth curve or total plant length (**Table 1**).

**Table 1.** . Aboveground plant traits were analyzed in l inear models using conditioning plant genotype, watering treatment as explanatory variables. P - values have been adjusted for multiple comparisons using the Benjamini - Hochberg correction. R ^2^m (marginal R ^2^) refers to the variance explained by fixed effects (i. e. watering treatment). In each model, experimental block was also included as a random effect (results not shown).

| Aboveground Plant trait | Watering Treatment | Inoculum (genotype of conditioning plant) | Inoculum*Watering treatment |
| --- | --- | --- | --- |
| **AUC Height** | p > 0.05<br>F=0.0042<br>df =1,29 | p > 0.05<br>$\chi^2=0.0890$<br>df =1 | p>0.05<br>$\chi^2=0.19$<br>df =1 |
| **Shoot Length** | **p.adj = 0.073**<br>F= 3.28<br>df = 1,101<br>R<sup>2</sup>m=0.018 | p>0.05<br>$\chi^2=0.37$<br>df =1 | p>0.05<br>$\chi^2=0.00$<br>df =1 |
| **Shoot weight (wet)** | **p.adj <0.0001**<br>F= 39.15<br>df =1,133<br>R<sup>2</sup>m=0.185 | p>0.05<br>$\chi^2=0.00$<br>df =1 | p>0.05<br>$\chi^2=0.00$<br>df =1 |
| **Shoot weight (dry)** | **p.adj=0.07**<br>F= 3.49<br>df =1,105<br>R<sup>2</sup>m=0.021 | p>0.05<br>$\chi^2=0.11965$<br>df =1 | p>0.05<br>$\chi^2=0.00$<br>df =1 |
| **Total plant weight (wet)** | **p.adj <0.0001**<br>F= 31.64<br>df =1,106<br>R<sup>2</sup>m=0.101 | p>0.05<br>$\chi^2=1.017$<br>df =1 | p>0.05<br>$\chi^2=0.00$<br>df =1 |
| **Chlorophyll Content** | **p.adj= 0.02**<br>F= 7.66<br>df =1,34<br>R<sup>2</sup>m=0.0247 | p>0.05<br>$\chi^2=2.1$<br>df =1 | p>0.05<br>$\chi^2=0.00$<br>df =1 |
| **Root to Shoot Length Ratio** | **p.adj = 0.015**<br>F=7.97<br>df =1,133<br>R<sup>2</sup>m= 0.0449 | p > 0.05<br>$\chi^2= 0.0000$<br>df =1 | p>0.05<br>$\chi^2=0.00$<br>df =1 |
| **Root to Shoot weight ratio (wet)** | **p.adj = 0.064**<br>F= 4.2<br>df =1,47<br>R<sup>2</sup>m= 0.013 | p > 0.05<br>$\chi^2 = 0.0000$<br>df =1 | p>0.05<br>$\chi^2 = 2.139$<br>df =1 |

The conditioning genotype predicted none of the aboveground traits measured in this experiment (**Table 1**). Further, the interaction between conditioning plant genotype and watering treatment did not predict any of the above ground traits measured (**Table 1**). As conditioning plant genotype did not have a significant effect on any aboveground trait, we did not further investigate whether MAP of conditioning plant site of origin had any effect on aboveground traits.

### Belowground plant traits were influenced by both watering treatment and the genotype of conditioning plant

As observed for aboveground traits, current water availability had a strong impact on belowground traits. The watering treatment predicted maximum root length, root wet weight, total endophyte density, septate fungal density, metaxylem area (normalized by total root area), total root length, and maximum root diameter (**Table 2**). Plants that received a drought treatment had roots that were 10.8% longer maximum length than that of well-watered plants. Well-watered plant roots were 21.6% heavier than droughted plant roots; however, the roots were still fresh when this measurement was taken. Droughted plants had 21.6% greater total endophyte density, as well as 57.4% greater septate fungal density specifically. Droughted plants also had 7% greater metaxylem area (normalized by root cross sectioned area), compared to their well-watered counterparts. Well-watered plants had 10.2% greater maximum root diameter. Watering treatment did not predict AMF density, root cross-sectioned area, aerenchyma area (normalized by root cross-sectioned area), number of root tips, average and median root diameter, or root surface area (**Table 2**).

**Table 2.** . Belowground plant traits were analyzed in l inear models using conditioning plant genotype, watering treatment as explanatory variables. P - values have been adjusted for multiple comparisons using the Benjamini - Hochberg correction. R ^2^m (marginal R ^2^) refers to the variance explained by fixed effects (i. e. watering treatment) . The effect s ize, or proportion of variance explained by each statistically significant predictor, has been included on the r ightmost column. In each model, experimental block was also included as a random effect (results not shown).

| **Belowground Plant Trait** | **Watering Treatment** | **Inoculum (Genotype of conditioning plant)** | **Inoculum\* Watering treatment** | **Percentage of variance attributable to random effects** |
| --- | --- | --- | --- | --- |
| **Root Maximum Length** | **p.adj = 0.076**<br>F=3.4687<br>df =1,112<br>R<sup>2</sup>m= 0.0189 | p > 0.05<br>$\chi^2 = 0.0422$<br>df =1 | p>0.05<br>$\chi^2 = 0.00$<br>df =1 | Inocula: 1.15%<br>Watering * Inocula: 0.0000006 % |
| **Root weight (wet)** | **p.adj = 0.000133**<br>F= 19.925<br>df =1,107<br>R<sup>2</sup>m= 0.052 | p > 0.05<br>$\chi^2 = 2.085$<br>df =1 | p>0.05<br>$\chi^2 = 0.00$<br>df =1 | Inocula: 3.39%<br>Watering * Inocula: 0.000000014% |
| **Total Endophyte Density** | **p.adj = 0.06**<br>F= 4.21<br>df =1,100<br>R<sup>2</sup>m= 0.025 | **p.adj =0.07**<br>$\chi^2 = 3.88$<br>df =1 | p>0.05<br>$\chi^2 = 0.00$<br>df =1 | Inocula: 11.42 %<br>Watering * Inocula: 0.00% |
| **AMF Density** | p>0.05<br>F= 0.041<br>df =1,100<br>R<sup>2</sup>m= 0.00025 | **p.adj=0.06**<br>$\chi^2 = 2.085$<br>df =1 | p>0.05<br>$\chi^2 = 0.00$<br>df =1 | Inocula: 13.8%<br>Watering * Inocula: 0.00% |
| **Septate fungi Density** | **p.adj = 0.008**<br>F= 10.375<br>df =1,98<br>R<sup>2</sup>m= 0.063 | p > 0.05<br>$\chi^2 = 2.41$<br>df =1 | p>0.05<br>$\chi^2 = 0.00$<br>df =1 | Inocula: 12.19 %<br>Watering * Inocula: 0.00 % |
| **Root Cross-Sectioned Area** | p > 0.05<br>F= 0.272<br>df =1,51<br>R<sup>2</sup>m= 0.0008 | p > 0.05<br>$\chi^2 = 0.00$<br>df =1 | p>0.05<br>$\chi^2 = 2.28$<br>df =1 | - |
| **Aerenchyma: Root Area Ratio** | p>0.05<br>F= 1.67<br>df =1,29<br>R<sup>2</sup>m= 0.006 | **p.adj=0.08**<br>$\chi^2 = 2.93$<br>df =1 | **p.adj=0.0076**<br>$\chi^2 = 9.25$<br>df =1 | Inocula: 11.02%<br>Watering * Inocula: 10.9 % |
| **Metaxylem: Root Area Ratio** | **p.adj = 0.02**<br>F= 5.52<br>df =1,353<br>R<sup>2</sup>m= 0.0125 | **p.adj=0.07**<br>$\chi^2 = 3.59$<br>df =1 | p>0.05<br>$\chi^2 = 0.00$<br>df =1 | Inocula: 6.25 %<br>Watering* Inocula: 0.00 % |
| **Number of Root tips** | p>0.05<br>F= 0.75<br>df =1,24<br>R<sup>2</sup>m= 0.004 | **p.adj=0.005**<br>$\chi^2 = 7.96$<br>df =1 | p>0.05<br>$\chi^2 = 0.0297$<br>df =1 | Inocula: 37.11 %<br>Watering*Inocula: 1.17 % |
| **Average Root diameter** | p>0.05<br>F= 0.4<br>df =1,72<br>R<sup>2</sup>m= 0.0029 | p>0.05<br>$\chi^2 = 0.87$<br>df =1 | p>0.05<br>$\chi^2 = 0.00$<br>df =1 | - |
| **Median Root diameter** | p>0.05<br>F= 1.2711<br>df =1,118<br>R<sup>2</sup>m= 0.01 | p > 0.05<br>$\chi^2 = 0.00$<br>df =1 | p>0.05<br>$\chi^2 = 0.00$<br>df =1 | - |
| **Maximum Root diameter** | **p.adj = 0.02**<br>F= 6.19<br>df =1,76<br>R<sup>2</sup>m= 0.043 | p > 0.05<br>$\chi^2 = 0.0207$<br>df =1 | p>0.05<br>$\chi^2 = 0.00$<br>df =1 | Inocula: 1.14%<br>Watering*Inocula: 0.00% |
| **Root surface area** | p>0.05<br>F= 2.62<br>df =1,70<br>R<sup>2</sup>m= 0.012 | **p.adj = 0.02**<br>$\chi^2 = 6.54$<br>df =1 | p>0.05<br>$\chi^2 = 0.00$<br>df =1 | Inocula: 23.75%<br>Watering* Inocula: 0.00% |

The rhizosphere inoculum (i.e., the genotype of the conditioning plant) predicted several belowground traits, specifically total endophyte density, AMF density, standardized aerenchyma area, standardized metaxylem area, number of root tips, total root length, and root surface area (**Table 2**). Belowground traits that were not predicted by genotype of conditioning plant included root maximum length, root wet weight, septate fungal density, root cross-sectioned area, and average, median, and maximum root diameters (**Table 2**). Oomycete presence/absence was not predicted by any of the variables (p>0.05, χ^2^ =0.52, df=3).

### MAP of conditioning plant’s home site was not correlated with AMF density, root aerenchyma area, and metaxylem area

Inoculum (*i.e.,* the genotype of the conditioning plant) marginally explained the variation in AMF densities, as well as variation in aerenchyma and metaxylem area in roots (**Table 2**). Upon further investigation using a simple linear model on the 33 genotypes’ best linear unbiased predictors resulting from the original mixed-effects linear models, we found that MAP of a conditioning plant’s home site did not further explain the variation in these traits (**Figures 4 and 5).**

**Figure 5.**
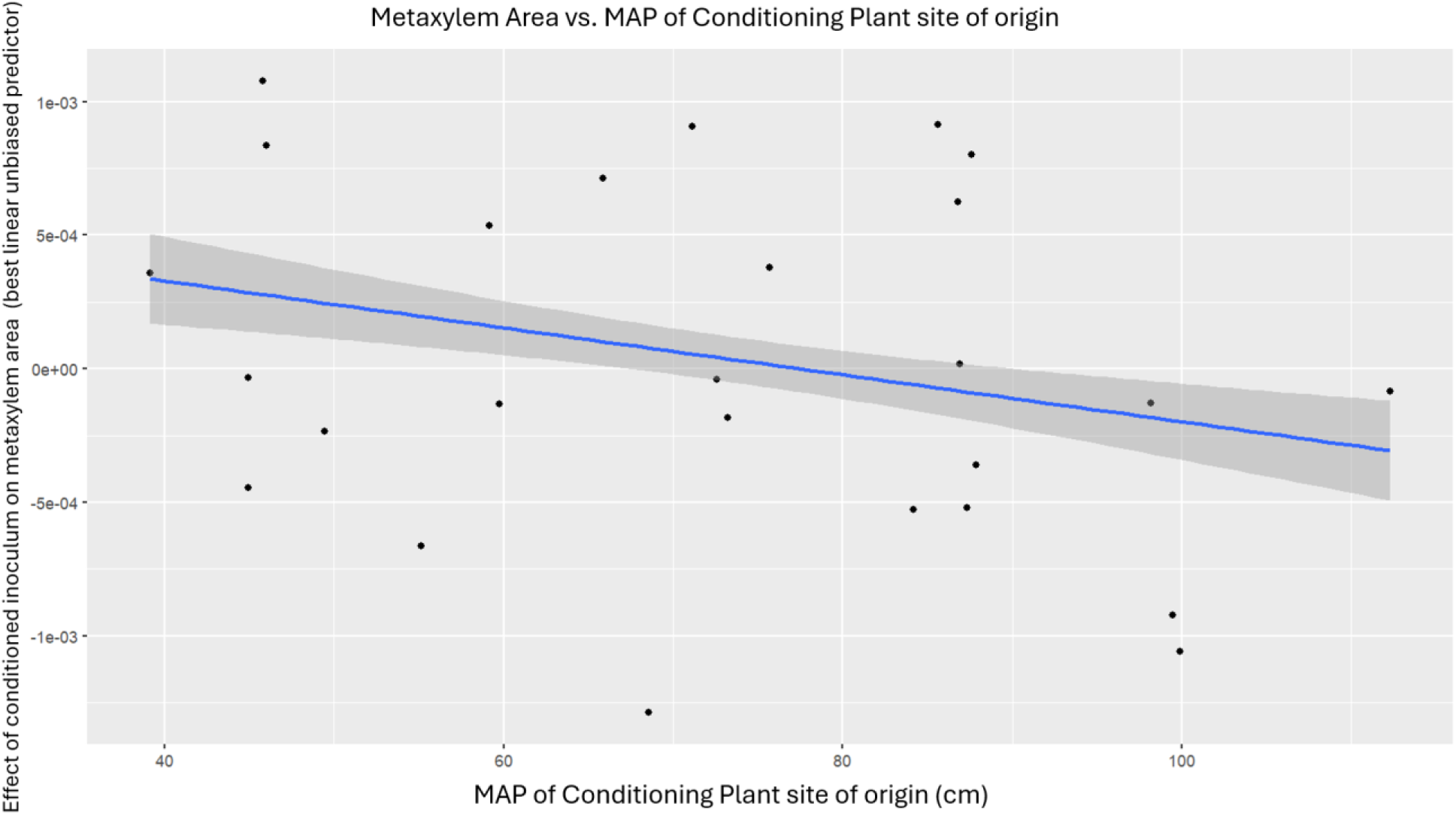
MAP of a conditioning plant site of origin was not correlated with metaxylem area (p > 0 . 05, F=0 . 127, df=1, 26, r = - 0 . 069).

### Properties of *T. dactyloides* root microbiomes

The compositions of the bacterial and fungal communities sequenced from the 33 rhizosphere inocula are comparable to previously sequenced rhizosphere communities in *T. dactyloides* (Kural-Rendon et al. 2023). The top ten bacterial classes were Actinobacteria, Alphaproteobacteria, Bacilli, Betaproteobacteria, Chitinophagia, Cytophagia, Deltaproteobacteria, Gammaproteobacteria, Sphingobacteria and Thermoleophilia (**Supplemental Figure 1**). The top ten bacterial families were Bacillaceae, Bradyrhizopiacaea, Burkholderiaceae, Comamonadaceae, Enterobacteriaceae, Oxalobacteraceae, Paenibacillaceae, Pseudomonadaceae, Sphingomonadaceae and Streptomycetaceae (**Supplemental Figure 2**). The top ten fungal orders were Chaetosphaeriales, Chaetothyriales, Cladosporiales, Eurotiales, Hypocreales, Mycosphaerellales, Phacidiales, Pleosporales, Sordariales and Xylariales (**Supplemental Figure 3**). The top ten fungal genera were Alternaria, Apodus, Cladosporium, Fusarium, Microdochium, Paraleptosphaeria, Rhinocladiella, Septoria, Setophoma, and Striaticonidum (**Supplemental Figure 4**). Many of these taxa were also found in previous work (Kural-Rendon et al. 2023). Many of the fungal genera sequenced were not annotated in the FungalTraits database (**Figure 6**); of the genera that were annotated, most were predicted to be soil saprotrophs, litter saprotrophs, or plant pathogens.

**Figure 6.**
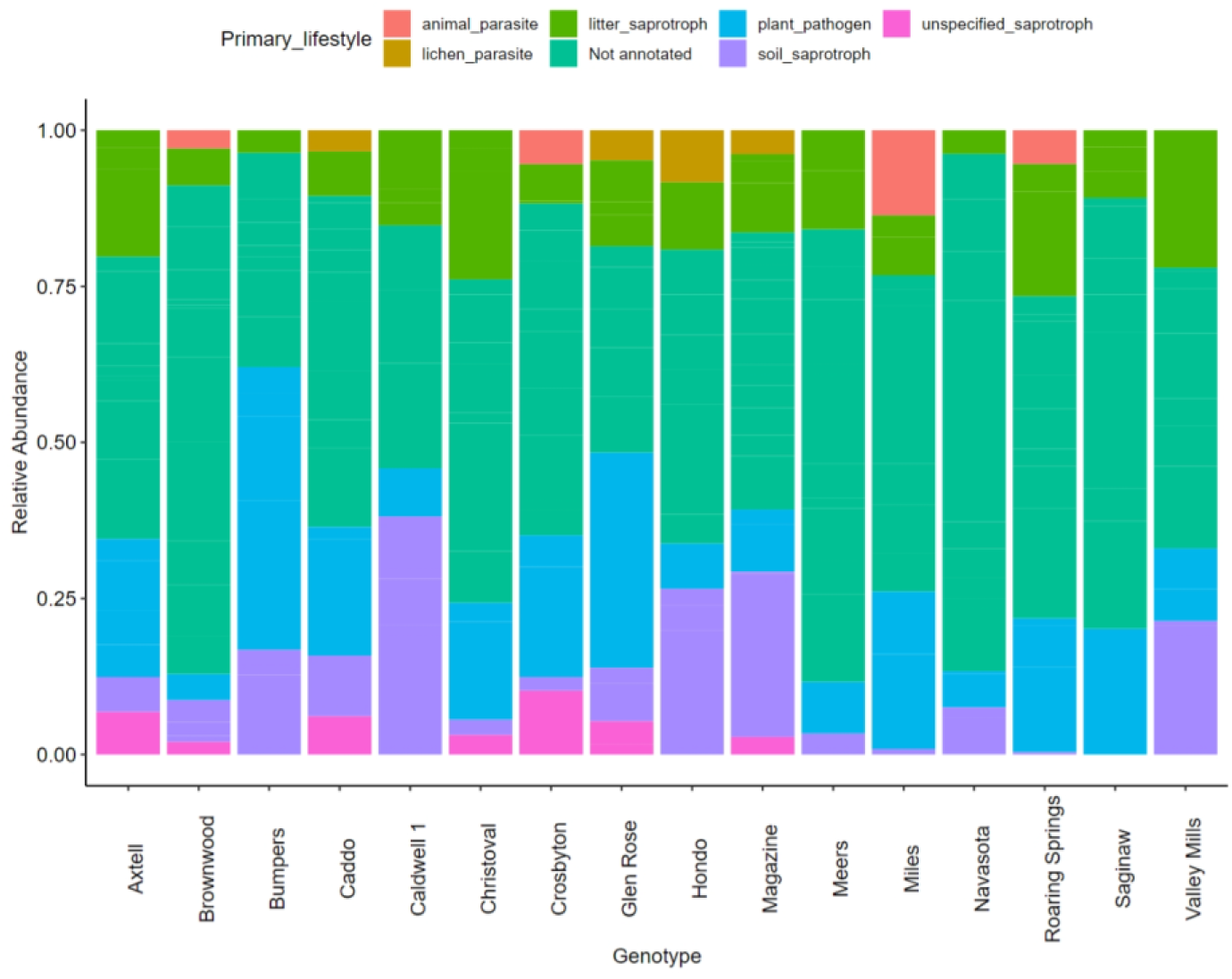
Primary functional groups of the fungal genera sequenced in our inoculum samples. The samples shown are what remained after quality control. Many of the genera sequenced in our samples were not annotated in the FungalTraits database.

Finally, we used Pearson correlation tests to determine how endophyte densities corresponded to plant traits both above and belowground. Aerenchyma and metaxylem areas, shoot length, root wet weight, number of root tips, total root length and root surface area all positively correlated with either total endophyte density, AMF or septate fungi specifically, or all three (**Table 4**). These traits increased as endophyte density increased. Root cross section area and average root diameter, however, negatively correlated with AMF density and total endophyte density, respectively (**Table 4**).

**Table 4.**
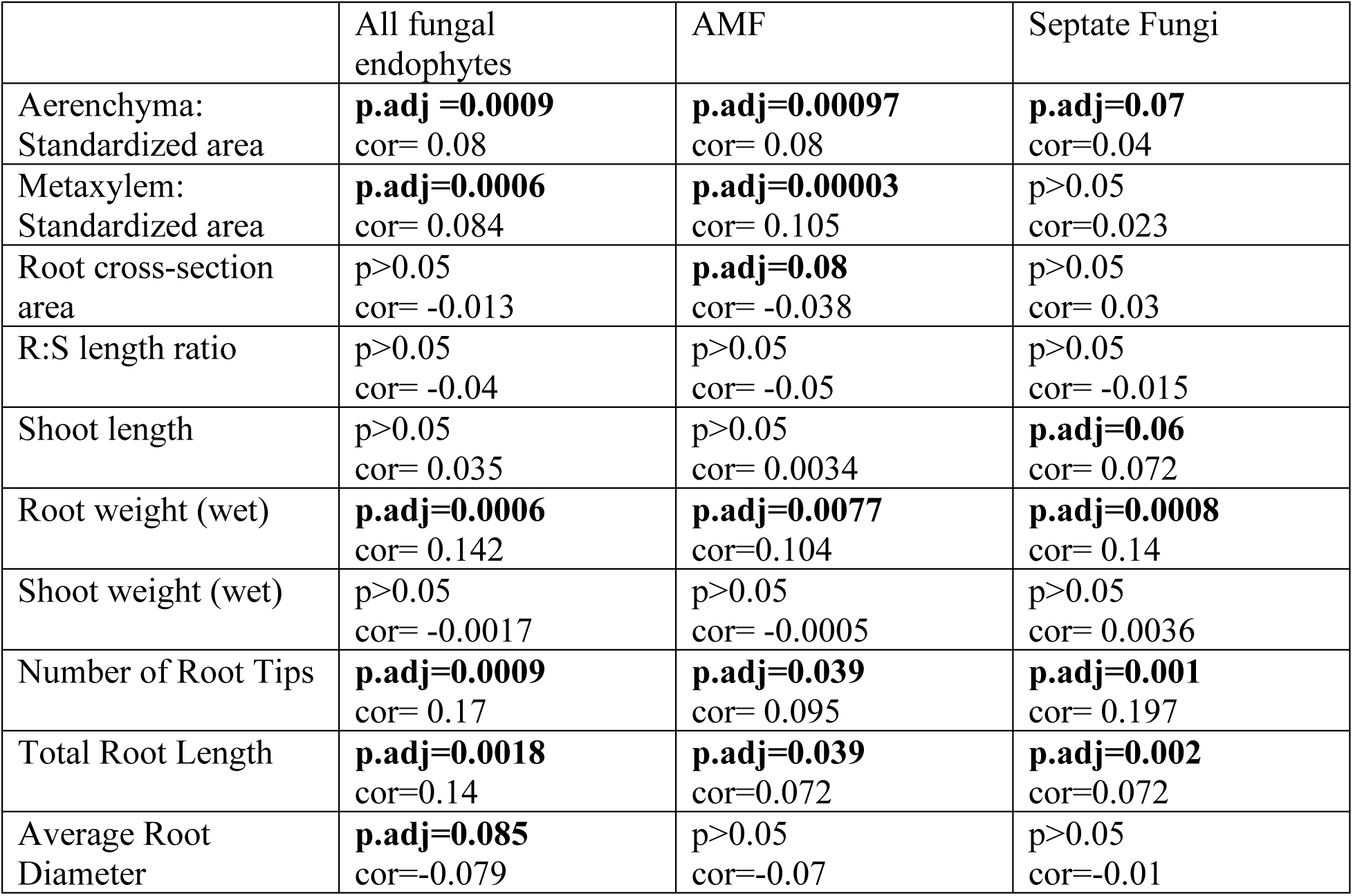
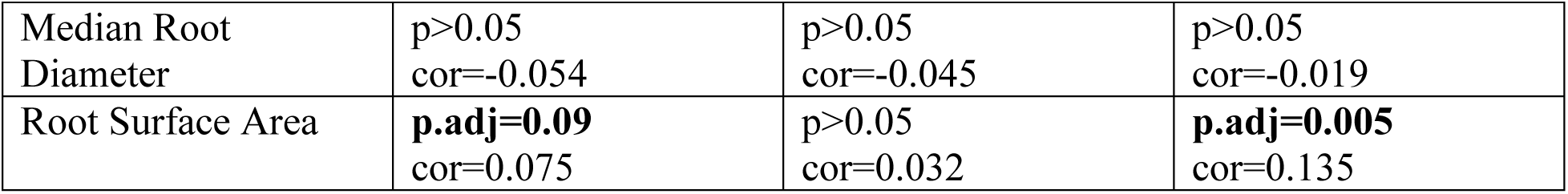
. Aboveground and belowground plant traits were analyzed in correlation tests against total endophyte, aseptate, and septate hyphal densities. P-values have been adjusted for multiple comparisons using the Benjamini - Hochberg correction.

|  | All fungal endophytes | AMF | Septate Fungi |
| --- | --- | --- | --- |
| Aerenchyma: Standardized area | **p.adj=0.0009**<br>cor= 0.08 | **p.adj=0.00097**<br>cor= 0.08 | **p.adj=0.07**<br>cor=0.04 |
| Metaxylem: Standardized area | **p.adj=0.0006**<br>cor= 0.084 | **p.adj=0.00003**<br>cor= 0.105 | $p>0.05$<br>cor=0.023 |
| Root cross-section area | $p>0.05$<br>cor= -0.013 | **p.adj=0.08**<br>cor= -0.038 | $p>0.05$<br>cor= 0.03 |
| R:S length ratio | $p>0.05$<br>cor= -0.04 | $p>0.05$<br>cor= -0.05 | $p>0.05$<br>cor= -0.015 |
| Shoot length | $p>0.05$<br>cor= 0.035 | $p>0.05$<br>cor= 0.0034 | **p.adj=0.06**<br>cor= 0.072 |
| Root weight (wet) | **p.adj=0.0006**<br>cor= 0.142 | **p.adj=0.0077**<br>cor=0.104 | **p.adj=0.0008**<br>cor= 0.14 |
| Shoot weight (wet) | $p>0.05$<br>cor= -0.0017 | $p>0.05$<br>cor= -0.0005 | $p>0.05$<br>cor= 0.0036 |
| Number of Root Tips | **p.adj=0.0009**<br>cor= 0.17 | **p.adj=0.039**<br>cor= 0.095 | **p.adj=0.001**<br>cor= 0.197 |
| Total Root Length | **p.adj=0.0018**<br>cor=0.14 | **p.adj=0.039**<br>cor=0.072 | **p.adj=0.002**<br>cor=0.072 |
| Average Root Diameter | **p.adj=0.085**<br>cor=-0.079 | $p>0.05$<br>cor=-0.07 | $p>0.05$<br>cor=-0.01 |
| Median Root Diameter | p>0.05<br>cor=-0.054 | p>0.05<br>cor=-0.045 | p>0.05<br>cor=-0.019 |
| Root Surface Area | **p.adj=0.09**<br>cor=0.075 | p>0.05<br>cor=0.032 | **p.adj=0.005**<br>cor=0.135 |

## Discussion

Our study investigated the extent to which historic precipitation explains the patterns of genetic variation that influence rhizosphere microbial recruitment in the wild grass *Tripsacum dactyloides*. In turn, rhizosphere microbial communities can affect the physiological traits of future conspecifics and how those conspecifics respond to differing precipitation regimes. While we found that the differing soil inocula and experimental watering treatment individually had various effects on different plant physiological traits, ranging from root architecture to chlorophyll content to absolute fungal abundance, we did not detect a significant interaction between historic environmental effects on soil microbial legacies and current experimental water regimes, except for aerenchyma area. While the research investigating intraspecific plant-soil feedbacks with respect to the conditioning plant’s environment of origin and its interaction with current watering treatment is limited, several studies have looked at the interplay between conditioning and experimental, or historic and current, watering treatments. A study by Kaisermann et al. in grasses found that previous drought led to changes in microbial communities that reduced the growth of subsequent conspecifics, and that plant responses to current watering treatment were dependent on historic watering treatment (2017). In contrast, another study showed that plant traits are minimally influenced by the interplay between historic and current watering, but that current drought does change the feedback between plants and their soil microbiomes (Enderle et al. 2024). Similarly, a study found that effect of historic conditioning did not play a significant role in plant performance under drought, suggesting that current watering conditions may play a more important role in plant performance (Wilschut and Van Kleunen 2021). The discrepancy in these findings may be due to the fact that plant-soil feedback is incredibly context dependent, and can shift depending on many factors such as plant species, soil type, environmental conditions – to name a few (Bennett and Klironomos 2019, Png et al. 2023), These conflicting results in the literature can also be due to differences in experimental design such as biotic factors (plant traits, microbial community composition), abiotic factors (soil type, drought severity, prior land use), and methodological differences (conditioning phase length and experimental phase length).

As expected, many aboveground and belowground traits were influenced by the watering treatment. A higher root to shoot ratio in drought treated plants compared to well-watered plants is consistent with the finding that plants will devote greater resources belowground during a drought period (Xu et al. 2015). A smaller root diameter is a common trait in drought-stressed plants (Comas et al. 2013), which are similar results to what we found in our drought treated plants.

None of the aboveground traits measured in this experiment were affected by the inocula. Several belowground traits measured were affected by the genotype of the conditioning plant. The difference in inocula effect on aboveground vs. belowground traits can be due to several reasons, primarily involving the direct contact between the soil microbes and the root structures. Microbes present in soil inocula directly colonize roots, leading to changes in root traits and morphology. Specifically, plant-growth promoting bacteria *Azospirillum* has been found to positively change root traits in ways that benefit plant host water nutrient uptake (Grover et al. 2021). Similarly, inoculation with beneficial *Bacillus* species in soybean plants have shown to impact roots by increasing root volume, length and branching – without affecting any aboveground traits (Araujo et al. 2021), perhaps due to resource allocation. Additionally, it is possible that inocula impact root traits more quickly than shoot traits, and if we had extended the ten week time frame of this experiment would have seen an effect on aboveground traits, as suggested by Koyama et al (2021).

Gao et al. show that belowground traits such as root length, density, as well as AMF fungal colonization can be shaped by different types of soil inocula (Gao et al. 2023). For a few traits where variation was explained by inoculum - AMF density, aerenchyma and metaxylem area - we found a specific effect of the MAP of conditioning plant site of origin. Aerenchyma area (normalized by cortex area) specifically was found be higher in plants adapted to drought conditions (Yamauchi et al. 2021b). This was the only trait in which we found that watering treatment and inoculum as well as the interaction between the two, account for a significant amount of variation in the trait. Similarly, increased metaxylem has been found to help plants improve water uptake efficiency in drought conditions (Kalra et al. 2024).

The role of AMF in conferring stress-including drought -tolerance to host plants has been established (Begum et al. 2019). Previous research in *T. dactyloides* has shown that plants from drier regions will have higher densities of fungal endophytes than plants from wetter regions (Kural-Rendon et al. 2023). A study similarly found increased AMF spore densities in a legume in the dry season, compared to the wet season ((Escudero and Mendoza 2005). In contrast, Zhang et al. found AMF hyphal density was positively correlated with precipitation in dominant forb species across an alpine steppe (Zhang et al. 2016). Other research in bilberry and a Chilean climbing vine has found a difference in fungal community diversity and composition along water availability gradients ((Guevara-Araya et al. 2022, Nguyen et al. 2024). Similar to plant soil feedback, AMF symbiosis with host plants is incredibly context dependent and can vary depending on many biotic and abiotic factors (Hoeksema et al. 2010). While all the experimental plants measured in this study shared a genotype (the “Pete” cultivar), the soil inocula that were used in our experiment were conditioned by 33 genotypes from widespread populations (Figure 1), which we initially assumed were adapted to different levels of precipitation, which in turn would have influenced the amounts of fungal propagules available to the experimental plants through genotype-specific rhizosphere effects. We hypothesized that this, in turn, would influence the experimental plants’ differential recruitment of fungal endophytes. However, our analysis did not show a specific influence of conditioning plant genotype home site precipitation on total endophyte and AMF densities, aerenchyma and metaxylem area ratios, or number of root tips or root surface area. This suggests that there could be other environmental factors of each conditioning plant’s site driving the differences we observed in the experimental plants’ traits, such as nitrogen availability (Kabir et al. 1997, Wang et al. 2017). We also found higher densities of fungal endophytes in roots with greater biomass. Due to the correlative nature of our analysis of the association between larger root cross sectioned area/greater root weight and fungal endophyte density, we cannot say whether our larger roots were large because they had more endophytes (*e.g.,* due to some growth-promoting effect of the fungi), or they had more endophytes because they were larger. A previous study conducted in both grass and forb species also found an increase in fungal endophyte biomass with an increase in plant belowground biomass (Eisenhauer et al. 2017). Similarly, we also found several root traits correlated with endophyte density, such as aerenchyma area, metaxylem area, number of root tips, total root length, average root diameter, and root surface area. AMF specifically can shape several root traits to increase the area it can colonize (Bergmann et al. 2020). Future work to determine if this relationship is causal can include an experiment to examine the effects of three levels of fungal inoculation (high, low, and no fungal inoculation), on root growth. This can perhaps be done in a split root experiment, to reduce individual plant variation in root growth.

Findings from this study can be useful in restoration, specifically in grassland and prairie systems. Plant genotype is an important consideration when working on a restoration project, specifically when using seeds or root stock. Previous work in tallgrass prairie grasses have demonstrated benefit from using both local and non-local genotypes in a restoration project, to increase genetic diversity, as well as instill some adaptability in the face of shifting climate patterns (Wilson et al. 2016). Another study recommends sourcing seeds not only from local populations but also from regions that match the projected future climatic conditions of the restoration site (Breed et al. 2018). More information about the environmental conditions which certain genotypes are adapted to can allow restoration practitioners to choose genotypes that may fare well under future environmental conditions. Additionally, field inoculation studies are becoming more common as the symbiosis between plants and their associated microbes, especially AMF, is a topic of increasing interest. Field inoculation with whole soils have been used to aid in restoration efforts to increase native plant diversity and reduce the risk of non-native reestablishment (Duell et al. 2023). Whole soil inoculation has also been found to increase crop yield and plant uptake of soil nutrients in agricultural settings (Bender and van der Heijden 2015). The interaction between certain inocula and different watering or precipitation regimes will continue to be an important topic of research in years to come. The legacy of drought on plant productivity and physiology through plant-soil feedback remains after the drought has ended (Lozano et al. 2022). Our results indicate that the selection of genotypes for inclusion in restoration projects may have implications for the resilience of the restored community to drought, via the mechanism of rhizosphere conditioning. The interplay between historic precipitation effects on plant populations and their genetics and shifting current (and future) precipitation and how it affects plant physiological responses are imperative to understand in order to accurately predict and prepare for how climate change will affect plant communities.

## Supporting Information

Supplemental Figures 1 and 2 display the relative abundances of the top ten bacterial classes and families, respectively. Supplemental Figures 3 and 4 display the relative abundances of the top ten fungal orders and genera, respectively.

## Acknowledgements

We thank Jim Kiniry and Amber Williams at the USDA ARS Soil and Water Laboratory for allowing us to sample from their collection. We thank Jaide Hawkins for assistance with rhizosphere sample collection and Maggie Wilson (REU Summer 2022) for her help in data collection. We thank Shannel Swiader for her help processing roots for imaging. We thank the University of Kansas Medical Center – Genomics Core for generating the sequence data sets.

## Author Contributions

CKR and MRW collected samples from Temple, Texas. CKR designed and conducted the experiment. NEF conducted DNA extractions and PCR. NEF and CKR conducted the amplicon sequence data analysis. CKR and KH both processed and analyzed root samples. CKR analyzed plant phenotype data. All authors contributed to the data interpretation, writing, editing and final approval of the manuscript.

## Data Availability

The sequence data will be available on NCBI. The code and all other data will be uploaded to Zenodo.

