## Supplemental Figures 1-4 for "Intraspecific plant-soil feedbacks alter root traits in a perennial grass"

Supplementary Figures

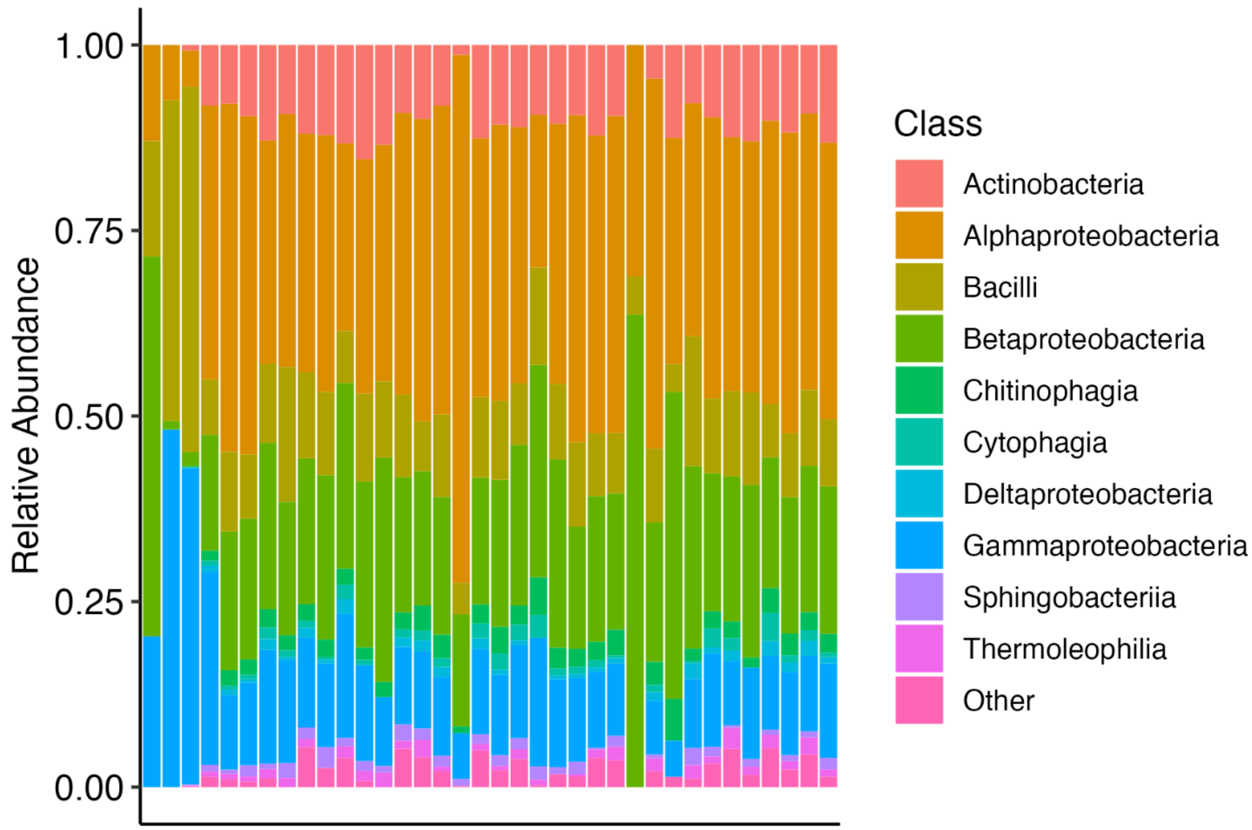


Supplemental Figure 1. Relative abundances of the top ten bacterial classes in the rhizosphere inocula.

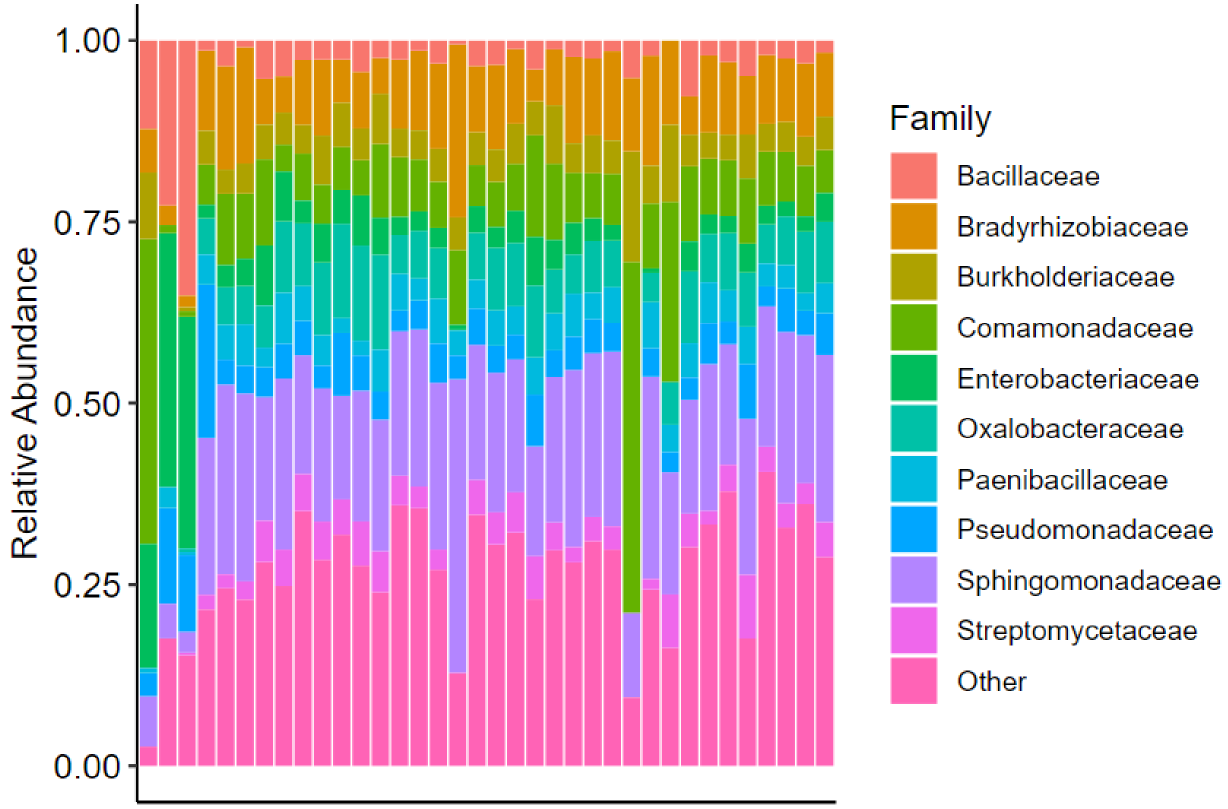


Supplemental figure 2. Relative abundances of the top ten bacterial families in the rhizosphere inocula.

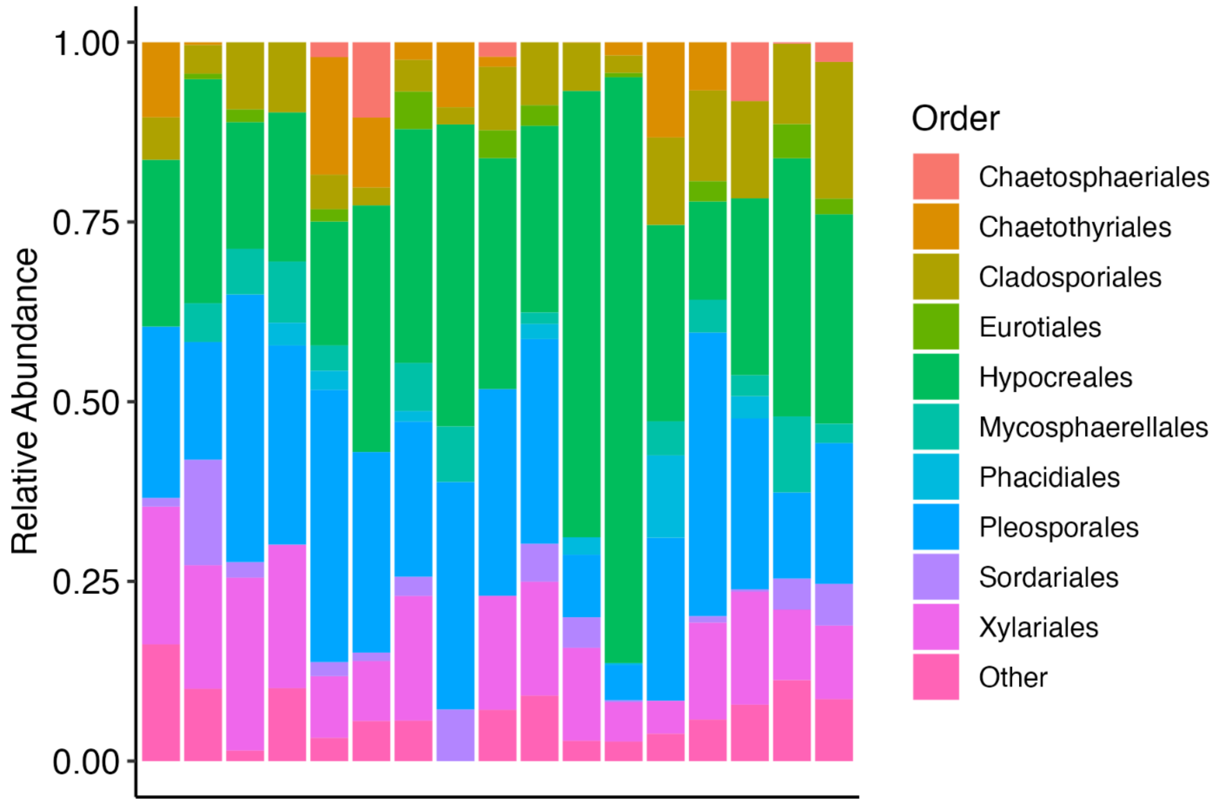


Supplemental Figure 3. Relative abundances of the top ten fungal orders in the rhizosphere inocula.

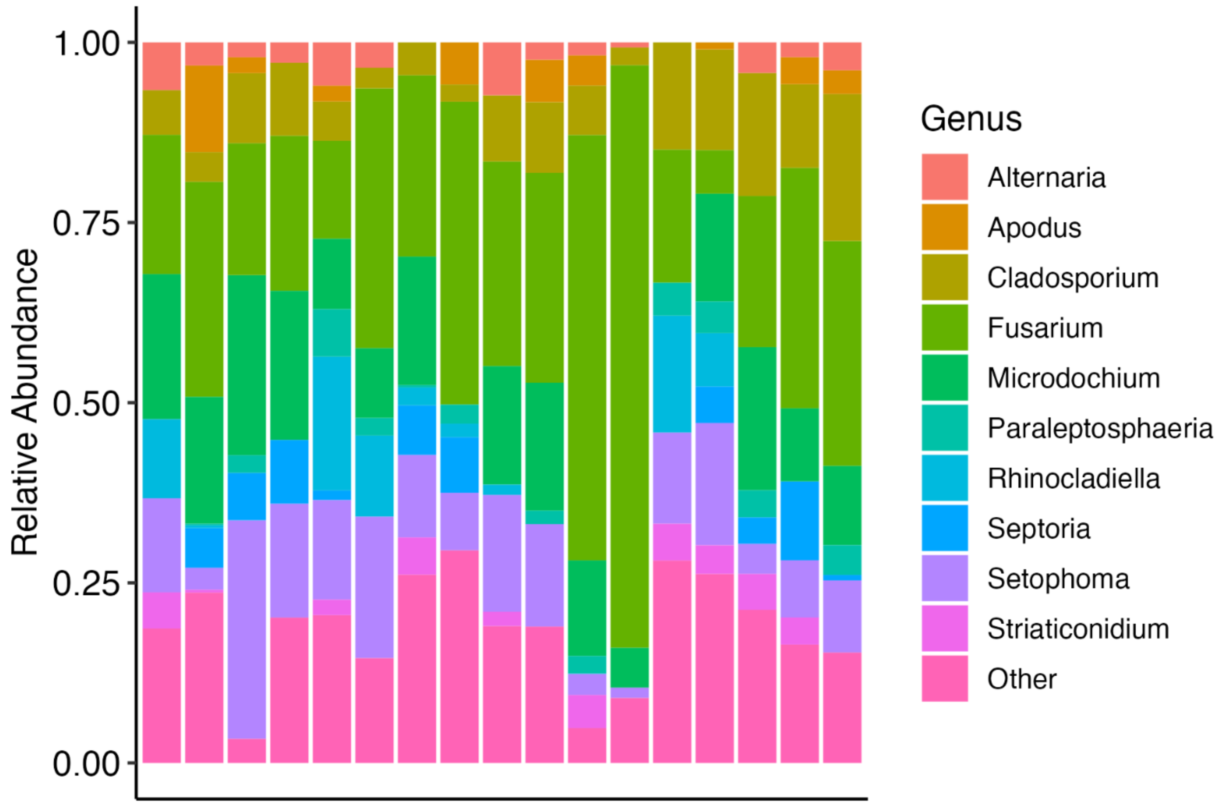


Supplemental figure 4. Relative abundances of the top ten fungal genera in the rhizosphere inocula.
